# Adipose-derived exosomal miR-421 targets CBX7 and promotes metastatic potential in ovarian cancer cells

**DOI:** 10.1101/2023.11.07.566022

**Authors:** Yi Zhang, Roslyn Tedja, Michael Millman, Terrence Wong, Alexandra Fox, Hussein Chehade, Meyer Gershater, Nicholas Adzibolosu, Radhika Gogoi, Matthew Anderson, Thomas Rutherford, Zhenggang Zhang, Michael Chopp, Gil Mor, Ayesha B. Alvero

**Author notes:** **Corresponding authors:** Ayesha B. Alvero, MD, MSc, C.S. Mott Center for Human Growth and Development Department of Obstetrics and Gynecology, Wayne State University 275 E. Hancock St., Detroit, MI 48201, Yi Zhang, PhD Neurology, Henry Ford Health System 2799 W Grand Blvd., Detroit, MI 48202. Authors contributed equally (Yi Zhang, Roslyn Tedja).

## Abstract

**Background:** Chromobox protein homolog 7 (CBX7), a member of the Polycomb repressor complex, is a potent epigenetic regulator and gene silencer. Our group has previously reported that CBX7 functions as a tumor suppressor in ovarian cancer cells and its loss accelerated formation of carcinomatosis and drove tumor progression in an ovarian cancer mouse model. The goal of this study is to identify specific signaling pathways in the ovarian tumor microenvironment that down-regulate CBX7. Given that adipocytes are an integral component of the peritoneal cavity and the ovarian tumor microenvironment, we hypothesize that the adipose microenvironment is an important regulator of CBX7 expression.

**Results:** Using conditioned media from human omental explants, we found that adipose-derived exosomes mediate CBX7 downregulation and enhance migratory potential of human ovarian cancer cells. Further, we identified adipose-derived exosomal miR-421 as a novel regulator of CBX7 expression and the main effector that downregulates CBX7.

**Conclusion:** In this study, we identified miR-421 as a specific signaling pathway in the ovarian tumor microenvironment that can downregulate CBX7 to induce epigenetic change in OC cells, which can drive disease progression. These findings suggest that targeting exosomal miR-421 may curtail ovarian cancer progression.

## Background

Cancer progression requires continuous acquisition of functions that support and increase aggressiveness and malignancy. Initial driver mutations, which can initiate malignant transformation are eventually followed by the creation of a pro-tumor microenvironment as cancer cells “educate” its surrounding stroma. In advanced stages, fibroblasts, immune cells, adipocytes, within this microenvironment would have been reprogrammed to primarily function as supporters of cancer growth and metastasis (1–3).

Adipose tissue is a central component of the cancer microenvironment (4), and is made up mostly of adipocytes and to a lesser extent, endothelial cells, and macrophages (5). Factors secreted by adipocytes as well as direct adipocyte-cancer cell interaction have been shown to support tumor progression in various cancer types including, breast, ovarian, prostate, and colon cancers (6–10).

Ovarian cancer (OC) is the most lethal of all gynecological malignancies and accounts for about 12,000 deaths annually (11). OC rarely metastasizes via the hematogeneous route, and almost exclusively metastasizes via shedding into the peritoneal cavity (12–14). Adipose-rich tissues within the abdomen are preferential sites for OC metastasis (15–18) (10, 17–19) and includes areas such as the omentum, mesosalphingeal/mesoovarian adipose, and adipose niches in the mesentery. Adipocytes have been shown to promote OC progression by providing fatty acids as an energy source to sustain rapid metastatic growth (20–22). In addition, adipocytes have been shown to enhance OC metastasis through secretion of chemotactic factors such as IL-8 and to promote chemoresistance by activating the Akt pathway (23) and upregulating the pro-survival protein Bclxl (24).

Chromobox protein 7 (CBX7) is a member of the Polycomb repressor complex 1 (PRC1). PRC1, and together with PRC2, are epigenetic regulators that induce histone modifications leading to repression of transcription (25–29). As part of PRC1, CBX7 is able to inhibit gene expression by binding to tri-methylated lysine 27 on histone 3 (H3K27me3) and inducing the monoubiquitination of lysine 119 on nearby histone 2A (H2Aub1) (30–33). CBX7 has been described as an oncogene, which is able to cooperate with c-Myc to induce aggressiveness in lymphomas (26) and contribute in the immortalization of fibroblasts (34). In other studies however, CBX7 has been described as a tumor suppressor and its downregulation has been correlated with aggressiveness and poor prognosis in thyroid, breast and colon cancers (35–38). Specifically, for OC, our previous work has demonstrated that the loss of CBX7 is correlated with enhanced metastatic potential and reduced patient survival (39). Mechanistically, CBX7 is able to achieve this by competing with Twist1 for binding to the E-box (39). As a result, Twist1, which is a master regulator of mesenchymal function cannot fully achieve its transcriptional role. Deletion of CBX7 using CRISPR/Cas9 relieves this inhibition allowing Twist1 to function and conferring enhanced metastatic potential to OC cells (39).

Exosomes are a subtype of small extracellular vesicles (EVs), which are released from cells into the extracellular space. They range in size from ∼30 to 150nm in diameter and carry various cargoes such as proteins, lipids, and nucleic acids, which reflect the composition of its cellular origin(40–42). In the tumor microenvironment, exosomes are secreted by cancer cells, infiltrating immune cells, endothelial cells, and cancer-associated fibroblasts (43–46), and mediate intercellular communication consequently contributing to various processes required for tumor progression such as cell proliferation, angiogenesis, and enhanced metastatic potential (43, 47–49).

Having demonstrated the significant impact of CBX7 loss on OC progression (39), we sought in this study to identify specific mechanisms within the ovarian tumor microenvironment that can regulate its expression. We report that adipose-derived exosomal microRNA-421 (miR-421) regulates CBX7 expression and function confers cancer cells with metastatic potential.

## Results

### Adipose-derived factors downregulate CBX7 and enhance migratory capacity in OC cells

Our group previously reported that CBX7 functions as a tumor suppressor in OC and its loss enhances metastasis in an OC xenograft model (39). In addition, *in vitro* migration through trans-well insert is also significantly augmented in OC cells upon CRISPR/Cas9-mediated knocked-out (KO) of CBX7 (Supp. Fig. 1). Thus, we hypothesize that factors secreted by the tumor microenvironment may be responsible for the regulation of CBX7 expression and function. Given the known significance of the adipose niche in OC progression, we posited that adipocytes may affect CBX7 expression. As such, we collected <u>a</u>dipose <u>c</u>onditioned <u>m</u>edia (ACM) from a human omental explant isolated from a female patient who underwent laparoscopic surgery for a benign gynecological condition and treated a panel of human OC cells. After 5 days of treatment, we observed that ACM are able to down-regulate CBX7 in all OC cells tested (Fig. 1A).

**Figure 1.**
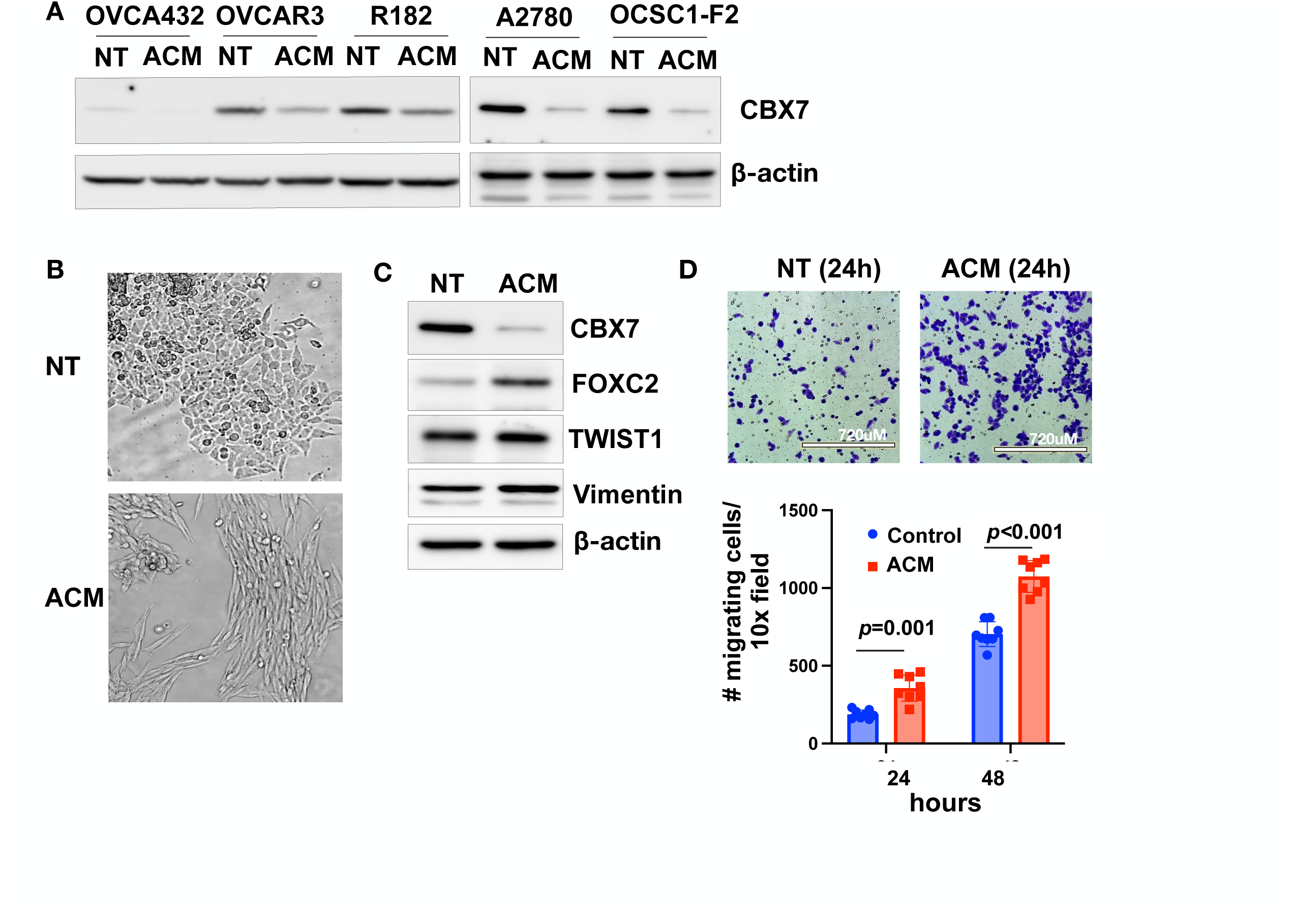
Adipose conditioned media (ACM) downregulate CBX7 and enhance migratory capacity in ovarian cancer cells. A panel of human ovarian cancer cells were cultured for 5 days in ACM from omental explants isolated from a female patient with a benign gynecological condition. No treatment control cells (NT) were cultured in growth media. **(A)** Levels of CBX7 were determined by Western blot analysis; **(B)** Representative microscopic images show acquisition of a fibroblastic spindle-like morphology; **(C)** Western blot analysis for EMT markers in OCSC1-F2 human ovarian cancer cells; **(D)** *top panel*, OCSC1-F2 human ovarian cancer cells were cultured in ACM or growth media for 5 days, trypsinized, washed, and placed in trans-well inserts with growth media. Note enhanced migration in cells previously cultured with ACM; *bottom panel*, quantification of migration (n=8); *** *p* < 0.0001. Data are presented as mean +/- SEM. Unpaired t-test was used to determine statistical significance.

Interestingly, ACM induced morphological changes characteristic of fibroblastic morphology (Fig.1B). Characterization of markers associated with epithelial mesenchymal transition (EMT) following treatment with ACM showed downregulation of CBX7, upregulation of the mesenchymal marker FOXC2, and maintained expression of Twist1 and Vimentin (Fig. 1C).

To evaluate if ACM is able to enhance functions that support metastatic potential, we collected OC cells after treatment with ACM and performed the trans-well migration assay. We observed that pre-treatment with ACM significantly enhanced *in vitro* migratory capacity compared to no treatment (Fig. 1D). Since the migration assay was performed in the absence of ACM, these results demonstrate that pre-treatment with ACM is able to confer cell autonomous mechanisms that can support metastatic potential. These results also show that ACM treatment recapitulates the enhanced migratory potential observed in CBX7 KO OC cells.

### Characterization of adipose-derived exosomes

Further characterization of CBX7 status after treatment with ACM showed that the down-regulation of CBX7 protein was not associated with a decrease in CBX7 mRNA (Supp. Fig. 2) suggesting a possible post-transcriptional mechanism. miRNAs are short non-coding RNAs that function in the post-transcriptional control of gene expression (50). miRNAs provide a specific sequence that recruits ribonucleoprotein (RNP) complex to the 3’ untranslated region (3’UTR) of target messenger RNAs (mRNAs) (51–53). The binding of miRNA-RNP complex to their target mRNAs results in mRNA degradation or translational repression and therefore downregulation of protein expression (54). We hypothesized that ACM downregulates CBX7 through its miRNA content. Since miRNAs are packaged and protected in EVs such as exosomes(55–57), we further hypothesized that these miRNAs may be carried in exosomes. Thus, we collected a new set of ACM obtained from 6 different patient omenta (Table 1) and performed differential ultracentrifugation to isolate exosomes. Characterization of the isolates with NanoSight showed homogeneous exosome preparations with average size of 100 nm (Fig. 2A). TEM analysis showed typical exosomal morphology of double layered cup-shaped membrane structure (Fig. 2B) and Western blot analysis showed the presence of tetraspanin markers, CD63 and CD9, and the absence of the endoplasmic reticulum marker, calnexin(58) (Fig. 2C) demonstrating that the collected particles are exosome-like small EVs.

**Figure 2.**
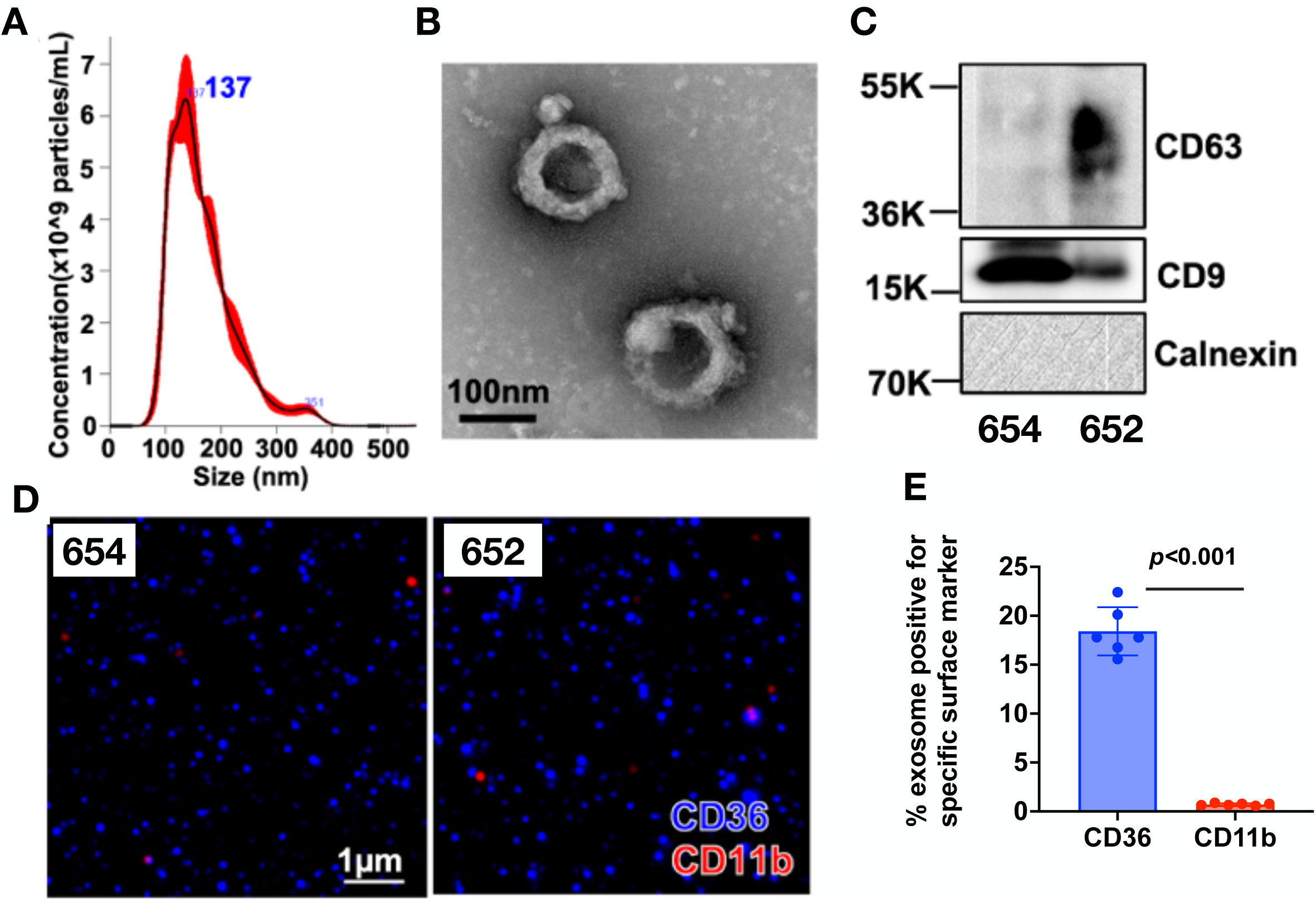
Characterization of exosomes derived from adipose conditioned media (ACM). Exosomes were isolated from ACM by differential ultracentrifugation. **(A)** Representative particle size distribution (ave = 137 nm) of ACM-derived exosomes analyzed by Nanoparticle Tracking Analysis; **(B)** Representative transmission electron microscopy image showing the doughnut-shaped morphology of the isolated exosomes; **(C)** Representative Western blot analysis of ACM-derived exosomes showing expression of exosomal markers CD63 and CD9 and absence of the endoplasmic reticulum marker, calnexin; Protein lysate of Adipose-Derived Mesenchymal Stem Cells (ADMSC) is used as positive control of calnexin. **(D)** Representative images from ExoView analysis showing majority of captured exosomes are positive for the adipocyte marker, CD36, and negative for the macrophage marker, CD11b; **(E)** Quantification of D and showing data from 6 individual patients. Data are presented as mean +/- SEM (n=6). Unpaired t-test was used to determine statistical significance. Data shown are for exosomes 654 and 652. Similar results were observed with other exosomal preparations.

**Table 1.**
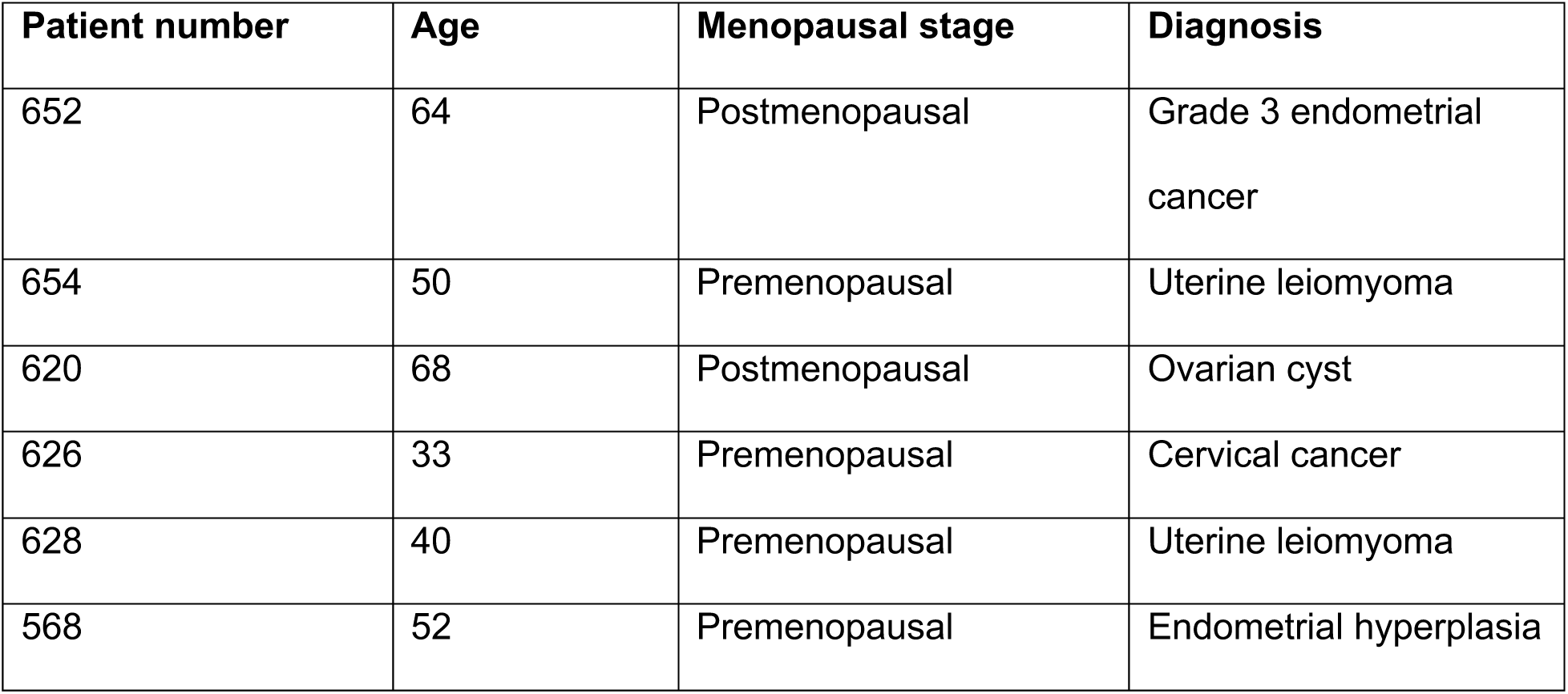
Patient characteristics.

The human omentum is primarily composed of adipocytes and immune cells within “milky spots” (59). To determine the cellular origin of the isolated exosomes, we utilized the ExoView platform to assay exosomal membrane proteins, which maintain the same characteristics as the cell of origin (60–62). Thus, we captured the exosomes using CD63, CD81 and CD9 antibodies and utilized CD36 and CD11b to identify the adipocyte and macrophage exosomal cell sources, respectively. Our results show significantly higher percentage of exosomes carrying the adipocyte marker, CD36 than the macrophage marker, CD11b (Fig. 2D and 2E), demonstrating that majority of the isolated exosomes from the ACM originate from adipocytes.

### Adipose-derived exosomes are sufficient to downregulate CBX7

To demonstrate that the isolated ACM-derived exosomes can be internalized by OC cells, we labelled the exosomes by their cargo RNAs with green fluorescence (GF-exosomes)(63, 64) and exposed them to mCherry-labeled OC cells. Microscopic analysis showed co-localization of GF-exosomes and mCherry OC cells, demonstrating the internalization of the labelled exosomes by the OC cells (Fig. 3A) and the putative delivery of their contents.

**Figure 3.**
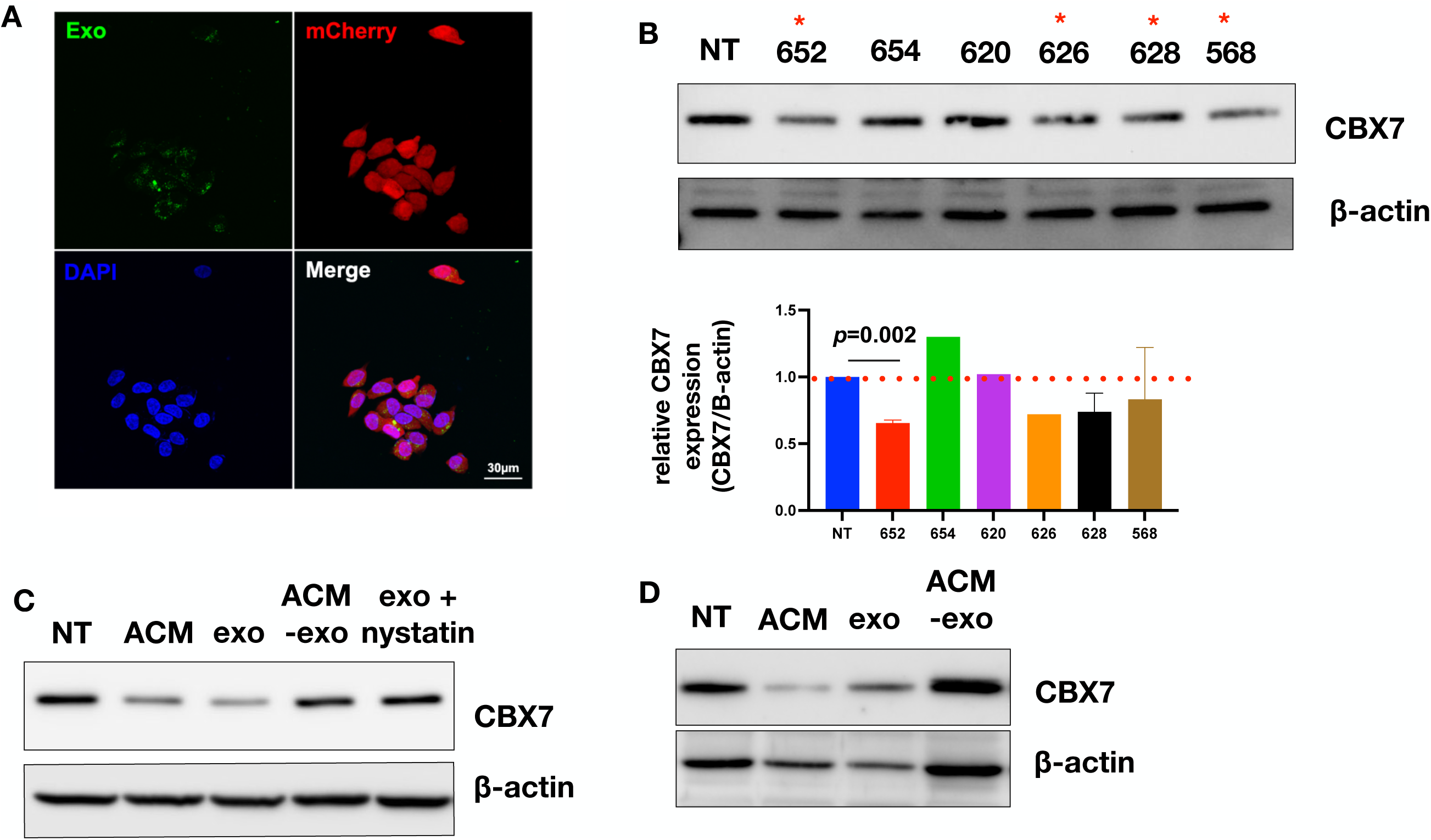
Adipose-derived exosomes are internalized by human ovarian cancer cells via endocytosis and induce down-regulation of CBX7. **(A)** Labelled ACM-derived exosomes were internalized by mCherry-labelled OCSC1-F2 cells; **(B)** Human R182 ovarian cancer cells were treated for 48h with adipose-derived exosomes (5×10^8^ exosomes/ml) isolated from patients in Table 1. *top panel*, levels of CBX7 determined by Western blot analysis. * denotes preparations able to down-regulate CBX7. *bottom panel*, densitometry quantification of top panel; **(C)** OCSC1-F2 human ovarian cancer cells were treated with ACM, exosomes (exo), exosome-depleted ACM (ACM-exo) or exosome with Nystatin and levels of CBX7 determined by Western blot analysis; **(D)** OVCAR3 human ovarian cancer cells were treated with ACM, exosomes (exo), or exosome-depleted ACM (ACM-exo) and levels of CBX7 determined by Western blot analysis.

We next determined the effect of ACM-derived exosomes on CBX7 expression. Interestingly, of the 6 exosome preparations tested, only 4 were able to down-regulate CBX7 (Fig. 3B). Patient 652, 626, 628 and 568 consistently down-regulated CBX7 compared to no treatment Control although only 652 reached statistical significance. To demonstrate that the effect on CBX7 is an exclusive property of the exosomes, we treated OC cells with ACM, isolated exosomes, or exosome-depleted ACM. Both ACM and isolated exosomes were able to decrease CBX7 expression, but this effect was lost in the exosome-depleted ACM (Fig. 3C and 3D). Finally, to further support that the observed decrease in CBX7 is due to exosome cargo, we treated OC cells with exosomes in the presence or absence of the endocytosis inhibitor, Nystatin. We observed that pre-treatment with Nystatin abrogated the effect of exosomes on CBX7 (Fig. 3C). Taken together, our results demonstrate that adipose-derived exosomes can down-regulate CBX7 in OC cells.

### Adipose-derived exosomal miR-421 targets CBX7 in OC cells

We next sought the determine the signaling pathway responsible for the observed changes in CBX7 expression. In keeping with the hypothesis that this may be secondary to exosomal miRNA cargo we utilized mirTarBase (https://mirtarbase.cuhk.edu.cn and TargetScanHuman (https://targetscan.org/) to generate a list of miRNAs predicted to target CBX7 (Table 2 and Table 3). We then cross-referenced this to published lists of exosomal miRNA that are known to be almost exclusively derived from adipocytes. The first list came from the comparison of circulating exosomal miRNA between AdicerKO mice (mice specifically lacking Dicer in adipose tissue generated using Cre-*lox* gene recombination) and wild-type controls (65). The second list came from the same study and a comparison of circulating exosomal miRNA between patients with congenital lipodystrophy (CGL) and healthy controls (65). Venn diagram analysis identified only one common miRNA from this list, which is miR-421 (Fig. 4A).

**Figure 4.**
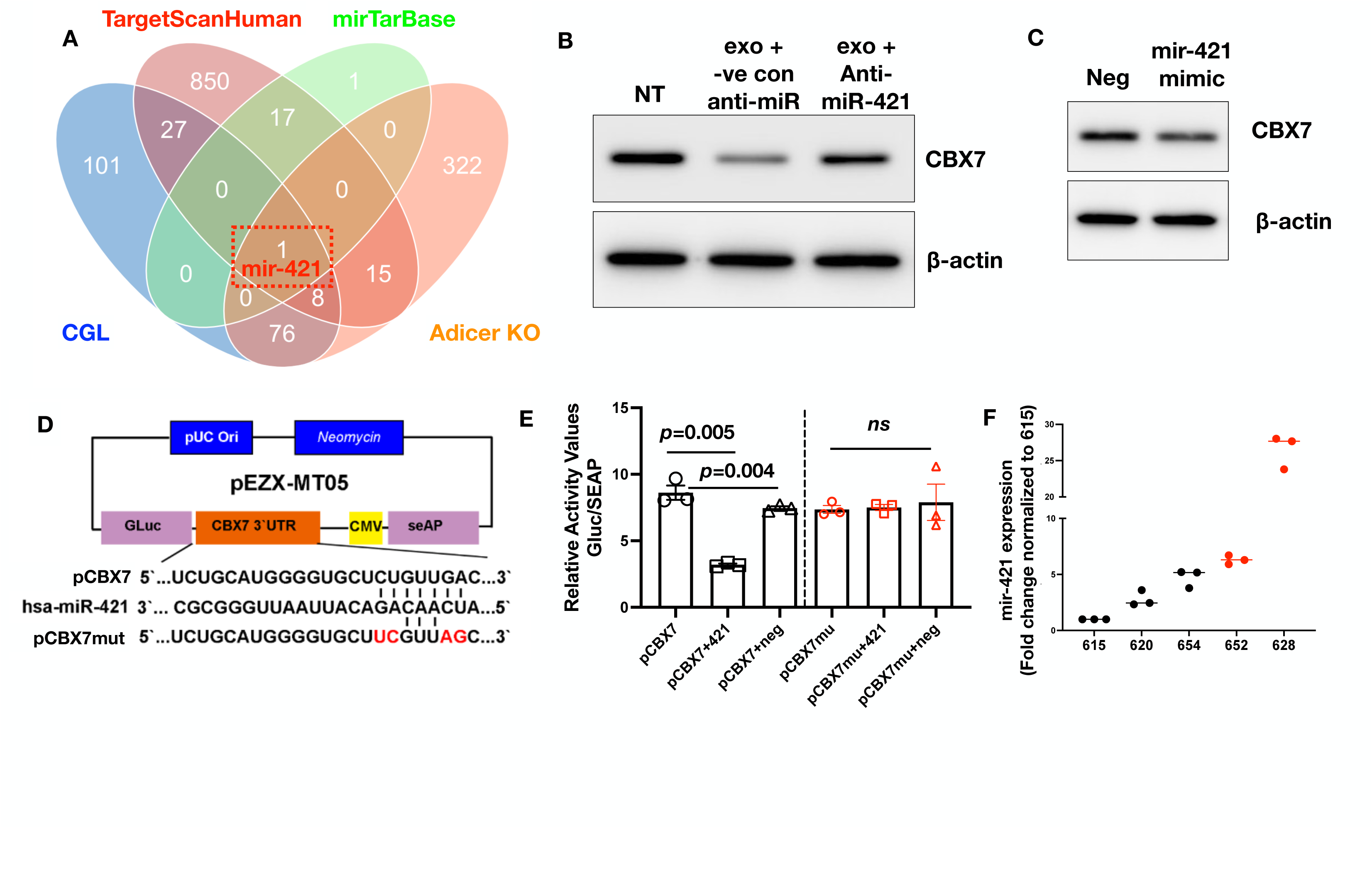
Adipose-derived exosomal miR-421 targets CBX7. **(A)** Venn diagram analysis from predicted miRNAs targeting CBX7 from mirTarBase (Table 2) and TargetScanHuman (Table 3) and lists of adipocyte specific miRNA from *Adicer* KO mice and patients with congenital lipodystrophy (CGL) (65) shows miR-421 as the only common element in the overlap; **(B)** OCSC1-F2 human ovarian cancer cells were treated with adipose-derived exosomes (exo) in the presence of anti-miR-421 or anti-miR negative control (-ve con Anti-mir) and effect on CBX7 was determined by Western blot analysis; NT, no treatment control **(C)** OCSC1-F2 human ovarian cancer cells were treated with miR-421 mimic and the effect on CBX7 was determined by Western blot analysis; Neg, non-specific miRNA; **(D)** plasmid design for luciferase reporter system carrying CBX7 3’ UTR predicted to be the binding site for miR-421 (pCBX7) and its mutated version (pCBX7mut). Sequence of human miR-421 (hsa-mir-421) is also shown; **(E)** Luciferase reporters were transfected in OCSC1-F2 human OC cells in the presence or absence of miR-421 or non-specific miRNA (Neg) as indicated, and levels of luciferase were measured as surrogate of binding affinity; **(F)** miR-421 was quantified in adipose-derived exosomes using qPCR. Note that exosomes successful in downregulating CBX7 (columns with red data points) contain higher levels of mir-421. Data are presented as mean +/- SEM (n=3). Ordinary One way ANOVA with Tukey’s multiple comparison test was used to determine statistical significance.

**Table 2.**
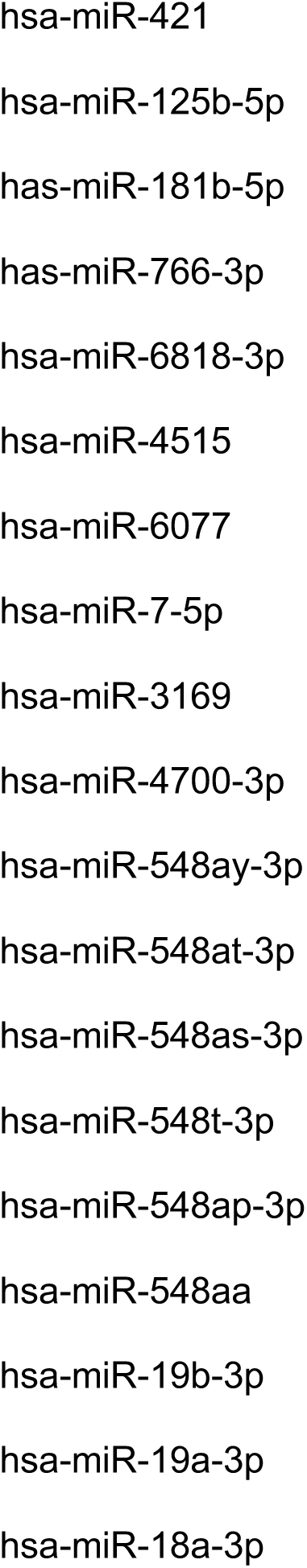
List of miRNAs predicted by mirTarBase to target CBX7.

**Table 3.**
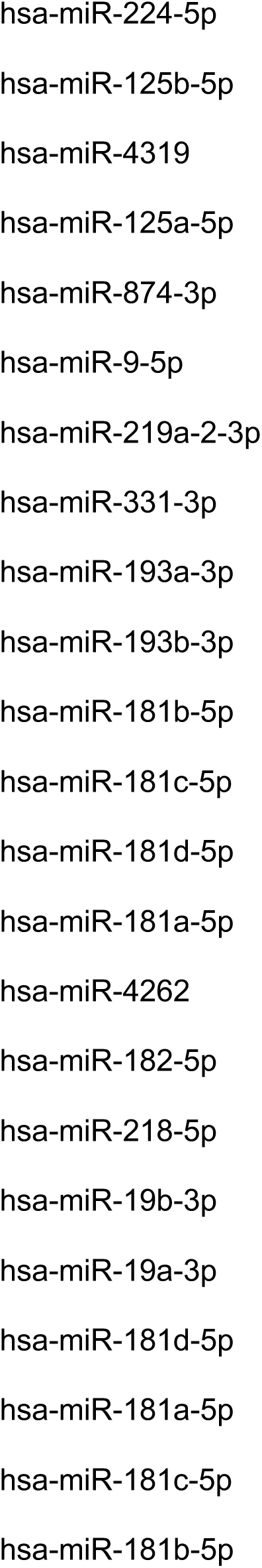
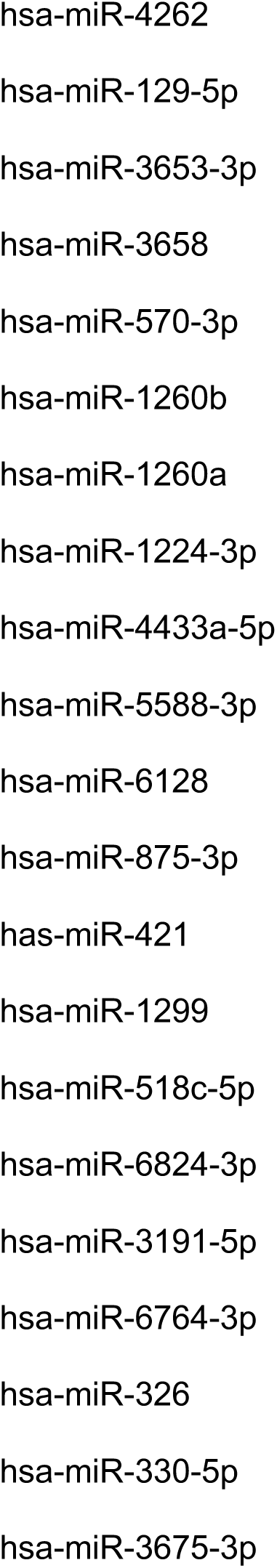
List of miRNAs predicted by TargetScanHuman to target CBX7 (45 out of 918 are listed)

We then proceeded to determine if miR-421 contributes to the exosomal signal that leads to ACM-induced CBX7 downregulation in OC cells. Thus, we treated OC cells with ACM derived exosomes in the presence or absence of anti-miR-421 or Control anti-miR. Our results show that the anti-miR-421, but not Control anti-miR, abrogated the effect of exosomes on CBX7 (Fig. 4B). In addition, we treated OC cells with miR-421 mimic and observed that transient expression of miR-421 mimic is sufficient to induce downregulation of CBX7 (Fig. 4C).

To conclusively demonstrate that CBX7 is a direct target of miR-421, we constructed a luciferase reporter upstream of CBX7’s 3’ UTR (pCBX7; Fig. 4D). Co-transfection of pCBX7 reporter in human OC cells with miR-421 resulted in a significant decrease in luciferase activity compared to pCBX7 reporter alone and pCBX7 co-transfected with Negative Control miRNA (Fig. 4E). This effect of miR-421 is specific to this sequence on CBX7’s 3’UTR as mutation of this sequence (pCBX7mut; Fig. 4D) abrogated the ability of miR-421 to decrease the luciferase signal (Fig. 4E). Collectively, these results demonstrate that the CBX7 gene is the direct target of miR-421 in OC cells.

### miR-421 is detectable in adipose-derived exosomes

In utilizing exosome preparations from 6 individual patients, we observed that only 4 of the 6 samples downregulated CBX7 (Fig. 3). Our final objective is to determine if we can detect miR-421 in the adipose-derived exosomes. We were able to detect mir-421 in our exosome preparations and in addition, we observed that exosomes that were able to induce CBX7 downregulation contained relatively increased levels of miR-421 (Fig. 4F). Taken together, our results show that miR-421 is a direct regulator of CBX7, and that adipose-derived exosomes are an important source of miR-421 in the ovarian tumor microenvironment.

## Discussion

We report a novel mechanism by which the adipose microenvironment can alter the phenotype of OC cells. In this study, we identified the existence of a miR-421/CBX7 axis that can lead to the downregulation of CBX7, a major component of the PRC. We demonstrate that the adipose niche is an important source of miR-421 and that it is packaged in exosomes. Our data demonstrate that adipose-derived exosomal miR-421 can function as an epigenetic regulator in OC cells by regulating CBX7.

Previously, we reported that CBX7 has tumor suppressor functions (39). Expression of CBX7 maintains OC epithelial phenotype by inhibiting mesenchymal triggers such as Twist1 (39).

Suppression of CBX7 in OC cells is associated with mesenchymal transformation and acquisition of metastatic potential. In this study we investigated the potential factors and sources responsible for the regulation of CBX7 expression and function, and consequently, the progression of OC. The findings reported here, highlight the contribution of the adipose niche in OC tumor progression.

CBX7 performs its gene-silencing function by modifying DNA-associated histones (25–29). As such, the genes repressed by CBX7, and hence the PRC complex, changes depending on cell state. As we previously reported, specifically for ovarian cancer, the expression of CBX7 alone is not predictive of survival (39). However, when used as a biomarker in conjunction Twist1, the absence of CBX7 in Twist1-positive ovarian cancer cells confers worse outcomes (ref). Whether or not CBX7 has a role in the initiation of ovarian cancer, such as exertion of pro-tumorigenic functions in normal ovarian surface epithelial cells or normal fallopian tube epithelial cells remains to be characterized. Nevertheless, our studies have shown that CBX7 is a major regulator of ovarian cancer progression (39) and that it can be regulated by factors secreted from adipocytes.

Secreted factors from the omentum have been shown to act as chemotactic factors that attract OC cells to this organ. Upon arrival, omental adipocytes have been shown to modulate OC cancer proliferation, metabolism, and response to chemotherapy(9, 66, 67). Given that OC mostly metastasizes via the trans-coelomic route and is mainly contained within the peritoneal cavity, the demonstration that adipose-derived exosomal miRNA can modulate epigenetic regulators in OC cells show that these cancer cells do not need to be in direct contact with adipocytes and can be located anywhere in the peritoneal cavity to be re-programmed by this microenvironment.

Although not fully characterized, selective loading of distinct molecules into exosomes is tightly regulated and does not occur randomly (68). Previous studies comparing intracellular versus exosomal miRNA content for instance, showed that some miRNAs are selectively packaged into exosomes, while specific miRNAs are actively excluded. This is true in both normal cells such as astrocytes and neurons (69) as well as cancer cells. In laryngeal squamous cell carcinoma for instance, miR-1246, miR-1290, miR-335-5p, miR-127-3p and miR-122-5p are selectively packaged in exosomes, while miR-4521, miR-4483, miR-30b-5p, miR-29b-3p and miR-374b-5p are selectively retained in the cell (70). A recent study has demonstrated that miRNA sequence motifs determine whether they are favorably sorted into exosomes or retained in cells(71). It has been postulated that this sorting may reflect the pro-survival benefits of the retained miRNAs to the cancer cells, while secreted miRNAs may provide benefit by regulating the tumor microenvironment. Still, this sorting mechanism is highly dynamic and can change depending on cellular state (72). It is therefore not surprising that not all ACMs tested were able to downregulate CBX7. It is tempting to speculate that pre-existing metabolic-associated risk factors known to significantly impact OC risk and survival such as body mass index(73), diabetes(74), or high-fat diet (75) can affect miRNA sorting into adipose-derived exosomes. Other conditions such as aging(76), menopause(77), and hormone replacement therapy(78) have been shown to affect the repertoire of circulating miRNAs. Moreover, the intracellular destination and processing pathways of exosomes after uptake by recipient cells are highly heterogeneous, which lead distinct biological activities in recipient cells (79). These correlations are currently under investigation in our lab.

Although free miRNAs can be detected in the circulation, the protective effect of exosomal membranes contribute to its stability (68). Indeed, exosomal miRNAs have been proposed as biomarkers for OC (80, 81). And, adipocyte derived exosomes have been successfully detected in blood (65). The demonstration that adipose-derived exosomal mir-421 can directly impact OC metastatic potential suggests the possible utility of exosomal miR-421 as a marker to identify patients needing closer follow up for recurrence.

## Conclusions

In conclusion, in this study we demonstrate that adipose-derived exosomal miR-421 within the tumor microenvironment downregulates CBX7 and consequently may enhance OC metastatic potential. These findings suggest that miR-421 is a molecular therapeutic target and a biomarker in OC patients.

## Materials and Methods

### Cell lines and culture conditions

OVCA432 (RRID:CVCL_3769) and OVCAR3 (RRID:CVCL_DH37) were obtained from ATCC (Manassas, VA) and cultured as instructed. The isolation and characterization of R182 and OCSC1-F2 have been previously reported (10, 82–85). These cells were cultured in RPMI1640 media supplemented with 10% FBS and 1% Pen/Step. Cells were grown in standard cell culture condition at 37°C with 5% CO2. For all experiments, cells were kept below 80% confluence. Mycoplasma testing was performed every 6 months and short tandem repeat (STR) profiling was done annually. Cells were used within 6 passages between experiments.

### Reagents and antibodies

Nystatin was purchased from MilliporeSigma (Cat. No. 1400-61-9). Mirvana miRNA mimic hsa-miR-421 (Cat. No. 4464066) was purchased from Thermo Scientific, Waltham, MA. miRIDIAN hsa-mir-421 hairpin inhibitor (Cat. No. IH-300996-03) was purchased from Horizon Discovery, (Waterbeach, UK). Lipofectamine RNAiMAX was purchased from Invitrogen (Waltham, MA).

Antibodies used were anti-CBX7 (RRID:AB_726005; 1:1000, Cat. No. ab21873, Abcam, Cambridge, UK), anti-β-actin (RRID:AB_2923704; 1:1000, Cat. No. 81115-1-RR, Proteintech, Rosemont, IL), anti-FOXC2 (RRID:AB_2798074;1:1000, Cat. No. 12974, Cell Signaling Technology, Danvers, MA), anti-Twist1 (RRID:AB_1130910; 1:500, Cat. No. CS-81417, Santa Cruz Biotechnology), anti-Vimentin (RRID:AB_10695459; 1:1000, Cat. No. 5741, Cell Signaling Technology, Danvers, MA), anti-GAPDH (RRID:AB_1078991; 1:10000, Cat. No. G8795, Sigma-Aldrich, St. Louis, MO).

### Human subjects and generation of adipose conditioned media

Sample collection was obtained with informed consent with prior approval from Karmanos Cancer Center and the University of South Florida IRB. Samples were consecutively collected from patients undergoing laparoscopic or open surgery for a benign or malignant gynecological condition irrespective of diagnosis or age. Upon receipt from the clinic, omentum tissues were minced with sterile razor blades and approximately 0.5 g of minced tissue was cultured per 100 mm tissue culture dish in 10 mL DMEM/F12 media supplemented with 1% exosome-depleted FBS (EXO-FBS-250A-1, System Biosciences, Palo Alto, CA). The following day, growth media were refreshed. and cell-free conditioned media were collected by centrifugation. Adipose tissue conditioned media (ACM) were stored at -80°C.

### Isolation of exosomes

Exosomes were isolated from the ACM via differential ultracentrifugation according to our published protocol (86, 87). Briefly, ACM was filtered through a 0.22 μm filter (MilliporeSigma, Burlington, MA) to remove all dead cells and large debris. The supernatant was then centrifugated at 10,000×g for 30 min followed by ultracentrifugation at 100,000×g (Optima XE-100 Ultracentrifuge, Beckman Coulter, Brea, CA) for 2 h. The resulting precipitate was re-suspended in phosphate-buffered saline (PBS) and used immediately or stored at 4°C.

### Characterization of exosomes

Collected particles were characterized according to MISEV 2018 guidelines (88). Briefly, the concentration and size distribution of exosomes were determined by Nanoparticle Tracking Analysis (NTA, NanoSight NSr300, Malvern, UK), according to our published protocol (86, 87). The ultrastructural morphology and the expression of exosomal markers were examined by transmission electron microscopy (TEM, JEOL, JEM 1400) and Western blotting, respectively (86, 87).

### ExoView single-EV profiling assay

After passing through 0.22 μm filter (MilliporeSigma, Burlington, MA), ACM were incubated with ExoView tetraspanin chips. Exosomes in ACM were captured by antibodies pre-coated on the chips, and against exosomal tetraspanin CD63, CD81 and CD9. A mouse IgG (MIgG) was used to capture non-exosomal particles. The captured exosomes were then triple-stained by fluorescent conjugated antibodies against the phenotypic marker of adipocyte (CD36) and macrophage (CD11b). Captured exosomes that were fluorescently positive for these proteins were counted and analyzed by ExoView analyzer. A fluorescent cutoff was set based on minimal detection of non-exosomal particles in MIgG capturing spots.

### Exosome labeling and internalization by OC cells

Exosomes were labeled using an RNASelect Green Fluorescent staining kit (Thermo Fisher) to stain exosomal cargo RNA(63, 64). Briefly, collected exosomes were incubated with 500nM labeling solution for 20 minutes at 37°C. After that, the labeled exosomes were rinsed in PBS and re-concentrated by ultracentrifugation. 5×10^8^ /mL labeled exosomes were then added into growth media of OCSC1-F2 human OC cells and incubated for 2h. The internalization of labeled exosomes (GF-exosomes) in F2 cells was then analyzed by green fluorescent confocal microscopy.

### Transwell Migration assay

1 × 10^5^ OCSC1-F2 human OC cells were resuspended in serum-free Opti-MEM (Thermo Scientific) and seeded on trans-well inserts with a polyethylene terephthalate membrane pore size of 8.0 μm (Cat. No. 353097, Corning, NY). The inserts were placed on 24-well cell culture plates with RPIM1640 media with 10% FBS. After 24 h, inserts were washed three times with PBS, fixed with 4% paraformaldehyde, permeabilized with 0.01% Triton X-100 (Sigma-Aldrich, St. Louise, MO), and stained with crystal violet Sigma-Aldrich, St. Louise, MO). Migrated cells were viewed and counted using a phase-contrast microscope (ECHO Revolve microscope, San Diego, CA).

### Luciferase reporter assay

The human 3’UTR of CBX7 gene encompassing the miR-421 binding site, and the 3‘UTR of *CBX7* gene with point mutations were cloned into a pEZX-MT05 vector with Gluc/SeAP duo-Luciferase reporter. The dual luciferase reporter system (Secrete-Pair ™ Dual Luminescence Assay, GeneCopoeia, MD) allowed the normalization of luminescence to minimize variations in transfection efficiencies and cell viability. Each vector was transfected in OCSC1-F2 cells by lipofectamine (Invitrogen, Waltham, MA) at a concentration of 2 ug vector/10^6^ cells. To assess the mRNA-miRNA interaction, each vector was co-transfected with miR-421 mimics (200pM/10^6^ cells, Cat. No. 4464066, ThermoFisher Scientific, Waltham, MA). Twenty-four hours later, the cells were lysed and luciferase activity was detected using a multimode microplate reader (PerkinElmer/Fusion, Waltham, MA).

### SDS-PAGE and Western blot analysis

Protein lysates were extracted from pelleted cells as previously described, and quantified using BCA assay (39, 89). Twenty micrograms of proteins were loaded in each well of 12% polyacrylamide gel and PAGE and Western blots were performed, as previously described (39, 89).

### RNA extraction and quantitative Reverse Transcriptase-Polymerase Chain Reaction (qRT-PCR) for CBX7

Total RNA was extracted from pelleted cells using RNeasy mini kit (Cat. No. 74106, Qiagen, Hilden, Germany). cDNA was obtained from 1 ug RNA using iScript cDNA synthesis kit (Bio-Rad, Hercules, CA). CBX7 and GAPDH qPCR was performed using SYBR Green Supermix (Bio-Rad, Hercules, CA) with 1:10 dilution of cDNA in a final volume of 10 μl according to manufacturer’s instructions. The following primer sequences were used for CBX7 (Forward primer: 5’-CGAGTATCTGGTGAAGTGGAAA-3’ and Reverse primer: 5’-GGGGGTCCAAGATGTGCT-3’) and GAPDH (Forward primer: 5’-TGACGCTGGGGCTGGCATT-3’ and Reverse primer: 5’-GGCTGGTGGTCCAGGGGT-3’). Primers were synthesized by Integrated DNA Technologies (San Diego, CA). qPCR was run on CFX96TM PCR detection system (Bio-Rad, Hercules, CA) using the following thermocycling parameters: 2 min 95°C; then 45 cycles of 20 s at 95°C and 1 min at 60°C. Control groups are incubated in growth media and experimental groups are incubated with ACM. No RT control was used as negative control. Relative expression was calculated using the comparative ΔΔCT method with Control group as reference. All reactions were performed in triplicates.

### RNA extraction and quantitative PCR for miR-421

Total RNA was extracted from ACM-derived exosomes using miRNeasy mini kit (Cat. No. 74106, Qiagen, Hilden, Germany). To determine miR-421 levels in exosomes, qPCR was performed with TaqMan miRNA assay kit (ThermoFisher Scientific, Waltham, MA). Briefly, miRNAs were reverse transcribed with the miRNA Reverse Transcription reagent and amplified with Taqman PCR reagents, which were specifically designed for detecting mature miR-421 sequence (AUCAACAGACAUUAAUUGGGCGC). U6 snRNA (mature sequence: GTGCTCGCTTCGGCAGCACATATACTAAAATTGGAACGATACAGAGAAGATTAGCATGGCCCCTGCGCAAGGATGACACGCAAATTCGTGAAGCGTTCCATATTTT) was used as the internal control. Relative expression was calculated by 2−ΔΔCt method according to published protocol (90).

### Statistical Analysis

Data were graphed and analyzed using Prism 9 (RRID:SCR_002798; GraphPad Software, Inc, Boston, MA). Unpaired t test or One-way ANOVA were used and p values <0.05 were considered statistically significant.

## Declarations

### Ethics approval and consent to participate

Sample collection was obtained with informed consent with prior approval from Karmanos Cancer Center and the University of South Florida IRB.

### Consent for publication

Not applicable.

### Availability of data and materials

Not applicable.

### Authors’ contributions

YZ – Conceptualization, Investigation, Methodology

RT – Investigation, Validation, Visualization

MM – Investigation

TW – Investigation

AF – Investigation HC – Investigation

MG – Investigation

NA – Formal analysis

RG – Resources

MA – Resources TR – Resources

ZZ – Conceptualization

MC – Conceptualization, Writing – review and editing

GM – Conceptualization, Funding acquisition, Writing – review and editing

AA – Conceptualization, Project administration, Investigation, Writing – original draft

---

**Supplementary Figure 1.**
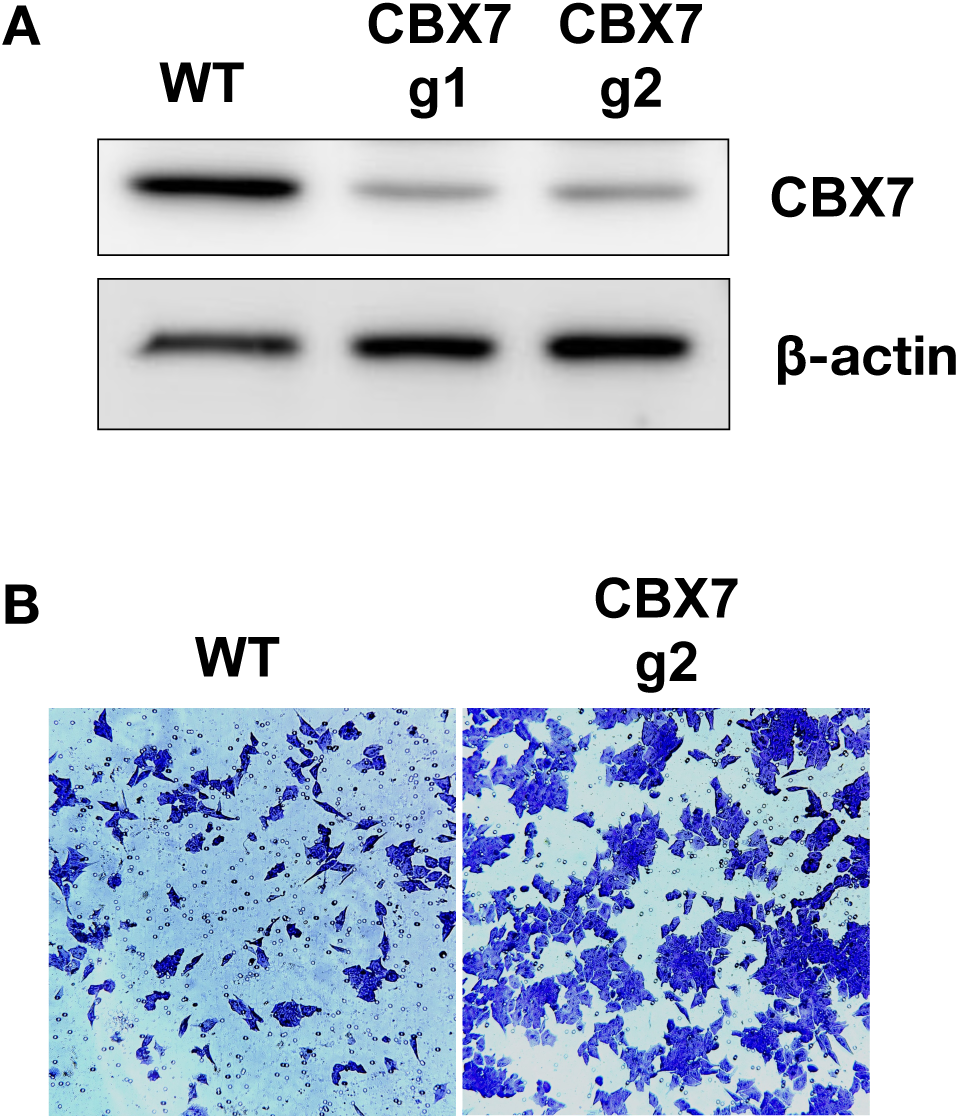
Loss of CBX7 is sufficient to enhance migration. **(A)** CBX7 was stably knocked-down in OCSC1-F2 human OC cells using CRISPR-Cas9. **(A)** Expression of CBX7 was determined by Western blot analysis; **(B)** Effect on migration was determined using trans-well migration assay. WT, wild-type; g1, guide RNA 1; g2, guide RNA 2.

**Supplementary Figure 2.**
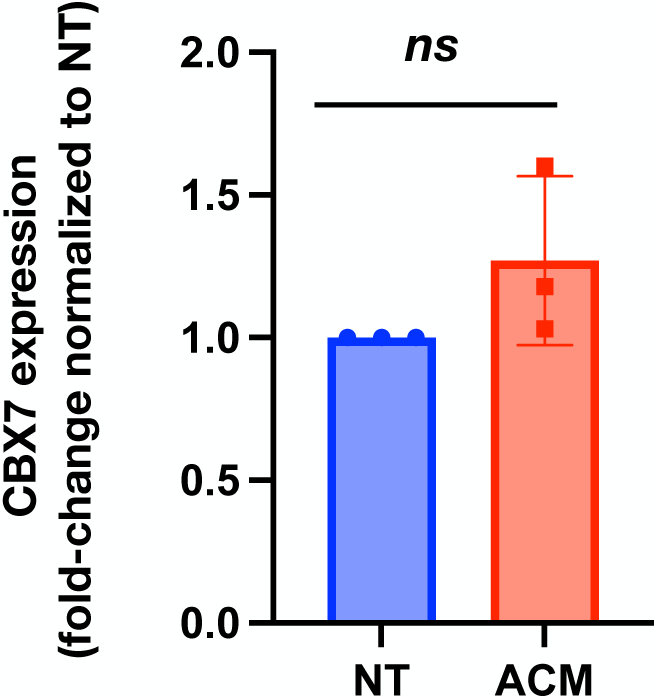
CBX7 protein downregulation is not associated with changes in mRNA. OVCAR3 and OCSC1-F1 human OC cells were cultured in ACM for 5 days and effect on CBX7 mRNA was determined by qPCR. NT, Control no treatment cells were cultured in growth media.

